# Epidermal Growth Factor Receptor Regulates Beclin-1 in Hyperoxic Acute Lung Injury

**DOI:** 10.1101/2024.05.24.595621

**Authors:** Zachary M. Harris, Asawari Korde, Johad Khoury, Edward P. Manning, Gail Stanley, Kennedy Mitchell, Ying Sun, Buqu Hu, Hyeon Jun Shin, John Joerns, Brian Clark, Lindsey Placek, Derya Unutmaz, Aigul Moldobaeva, Lokesh Sharma, Maor Sauler, Govindarajan Rajagopalan, Xuchen Zhang, He Wang, Mahboobe Ghaedi, Min-Jong Kang, Jonathan L. Koff

## Abstract

While delivery of supplemental oxygen is a life-saving therapy, exposure to high levels of oxygen, called hyperoxia, is associated with increased mortality in the intensive care unit (ICU). Hyperoxia leads to oxidant-mediated acute lung injury (ALI) and pulmonary cell death, called hyperoxic acute lung injury (HALI). Elucidation of molecular mechanisms in HALI could identify therapeutic targets in ALI. In the current study, we examined *in vivo* effects of HALI on Beclin-1 (BCN1), a molecule that regulates autophagy and cell death. Effects of HALI on BCN1 and autophagy markers were examined in wildtype mice. Analysis of BCN1 and autophagy was completed via Western blot, RT-qPCR, and immunohistochemistry. In wildtype mice, HALI led to increased BCN1 in the lung and in the alveolar epithelium. HALI resulted in significant alterations in markers of autophagy in the lung, including reduced microtubule-associated protein 1B-light chain (LC3B)-II/-I ratios, suggesting reduced autophagic flux. HALI caused increased LDH release in human alveolar type-II cells derived from induced pluripotent stem cells (AT2s^iPSC^), as well as reduced LC3B-II/-I ratios. We previously showed that inhibition of the tyrosine kinase receptor epidermal growth factor receptor (EGFR) is protective in HALI. EGFR^Wa5/+^ mice, which have genetically reduced EGFR activity and improved survival in HALI, showed increased total BCN1, reduced phosphorylated-(p-)/total BCN1 ratios, and decreased LC3B-II/-I ratios in the lung in HALI compared with wildtype. Administration of wortmannin, a phosphatidylinositol-3 kinase (PI3K) inhibitor which decreases BCN1-mediated autophagy, led to increased mortality in HALI in wildtype mice. These data support that regulation of BCN1 and autophagy by EGFR is a protective mechanism in HALI, a pathway which warrants further study for its therapeutic potential.

**KEY MESSAGES:** HALI is associated with increased mortality in the ICU and causes alveolar epithelial cell death, but the role of BCN1 in HALI is not well defined. This study shows that BCN1, a molecule involved in autophagy and cell death, is regulated by EGFR in HALI *in vivo*. These results are significant because regulation by BCN1 and autophagy by EGFR is a novel pathway in HALI with therapeutic potential that warrants further study.

## INTRODUCTION

More than 300,000 patients are placed on mechanical ventilation in the intensive care unit (ICU) annually in the United States (1–3). Many of these patients will require high oxygen, called *hyperoxia*, during their clinical course (4, 5). While oxygen is necessary to prevent tissue hypoxia, exposure to hyperoxia is common in the ICU and is associated with increased mortality (6–8).

Exposure to hyperoxia leads to a type of oxidant-mediated acute lung injury (ALI), called hyperoxic acute lung injury (HALI) (9). The pathogenesis of HALI involves epithelial cell death in the lung (10, 11). In patients with acute lung injury (ALI)/acute respiratory distress syndrome (ARDS) who require hyperoxia to maintain proper oxygenation, hyperoxia activates pathways that lead to cell death (10, 12, 13). It is critical to identify mechanisms of cell death in HALI to elucidate lung pathobiology for patients with ALI/ARDS who require high oxygen. In addition, HALI is a model for “sterile” (i.e., non-infectious) ARDS induced by oxidative stress. Despite the large number of patients hospitalized with ARDS each year and high mortality for this condition (14), there are no medical therapies for ARDS. Thus, elucidation of novel therapeutic targets for ARDS is critical. Identifying therapeutic targets in HALI has translational potential for therapeutic targets in ARDS.

Autophagy is a dynamic, homeostatic process in which stress conditions induce formation of autophagosomes that sequester unwanted contents for recycling (15, 16). While generally considered beneficial (17), the role of autophagy in HALI is not clear (18). Coiled-coil moesin-like BCL2-interacting protein [Beclin-1 (BCN1)] induces autophagy by forming a key signaling complex through highly regulated mechanisms. BCN1 contains a BCL2 homology 3 (BH3) domain, and because B-cell lymphoma 2 (BCL2) molecules contain and signal through BH1-4 domains (19), BCN1 acts as a nexus between apoptosis and autophagy (20). In addition, BCN1 contains distinct phosphorylation sites which, when activated, can either promote or downregulate autophagy (20). BCN1 signaling, which affects both autophagy and apoptosis through interrelated signaling pathways (20), makes BCN1 a notable therapeutic target in diseases involving regulated cell death (21), such as HALI (10, 11) and ALI/ARDS (22).

Epidermal growth factor receptor (EGFR) is a receptor tyrosine kinase with pleiotropic cellular effects that, when activated, is involved in multiple cellular processes that are beneficial in ALI. These include alveolar epithelial cell proliferation (23, 24), alveolar epithelial cell migration (23, 25, 26), epithelial cell barrier restoration after injury (27), wound repair in airway (28) and alveolar epithelial cells (26), and respiratory epithelial repair following naphthalene-induced ALI (29, 30). However, EGFR activation has also been shown to be deleterious in ALI, such as in ventilator-induced lung injury (VILI) (31) and lipopolysaccharide (LPS)-induced ALI (32). In addition, EGFR activation can be deleterious in asthma (33), pulmonary fibrosis (34, 35), and viral infection (36–38).

We previously showed that mice with genetically reduced EGFR activity have improved survival, reduced ALI, and decreased type-II alveolar epithelial apoptotic cell death in HALI (39). In addition, we previously showed that pharmacologic inhibition of EGFR led to reduced apoptotic cell death in type-II alveolar epithelial cells *in vitro* (39). Based on the pleiotropic effects of EGFR, inhibition of EGFR is likey beneficial in HALI via additional mechanisms. EGFR regulates BCN1 in cancer, which affects autophagy (40–42). To our knowledge, EGFR regulation of BCN1 has not been studied in non-cancer cells. In addition, EGFR regulation of BCN1 has not been studied in HALI. Based on this, we hypothesized that regulation of BCN1 and autophagy provides a mechanism in which EGFR inhibition provides protection in HALI.

In this study, we examined the effects HALI on BCN1 and autophagy markers in mice and in human alveolar epithelial type-II cells derived from induced pluripotent stem cells (AT2s^iPSC^). *In vivo*, we found that BCN1 is increased in HALI in the lung and in the alveolar epithelial compartment. HALI led to significant alterations in autophagy markers in the lung *in vivo*, suggesting decreased autophagic flux. Genetically altered mice with reduced EGFR activity (EGFR^Wa5/+^ mice), which we have previously shown to have improved survival and reduced alveolar epithelial apoptosis in HALI (39), contained increased BCN1 and decreased phosphorylated-(p-)/total BCN1 ratios in the lung in HALI. In HALI, EGFR^Wa5/+^ mice had decreased LC3II/-I ratios, suggesting reduced autophagic flux. In addition, administration of the phosphatidylinositol-3 kinase (PI3K) inhibitor wortmannin led to increased mortality in HALI *in vivo*. These data implicate that BCN1 and autophagy are regulated in HALI and modulated by EGFR. BCN1-mediated autophagy and its regulation via EGFR is a novel cell death pathway in HALI with therapeutic potential in oxidant-mediated ALI that warrants further study.

## MATERIALS AND METHODS

### Mice

EGFR-deficient mice containing the Wa5 Egfr allele (EGFR^Wa5/+^ mice) were a generous gift from Dr. David Threadgill at Texas A&M College of Medicine. The Egfr Wa5 allele contains a missense mutation within the EGFR coding region that results in decreased EGFR activity (43). Thus, Egfr Wa5 functions as a hypomorphic Egfr allele. The genetic background of EGFR^Wa5/+^ mice is BALB/c, C3H, and C57Bl/6J (43). Mice were kept and maintained in the Yale Animal Resource Center.

### Oxygen Exposure

Adult eight-to twelve-week-old mice were exposed to 100% oxygen delivered continuously in a plexiglass hyperoxia chamber as previously described (44). For survival experiments, mice were closely monitored, and time of death was recorded. To inhibit the BCN1-vacuolar protein sorting 34 (VPS34) complex, the mold metabolite wortmannin (Selleck, Houston, TX, USA; catalog # S2758) was used (45). Wortmannin (1 mg/kg) and vehicle control were administered via intraperitoneal (I.P.) injection immediately prior to initiation of oxygen exposure and subsequently every 24 hours for a total of three injections (i.e., mice were injected at initiation of oxygen exposure, 24 hours, and 48 hours). For timed exposures, mice (n=4-6 per group) were exposed to 100% oxygen continuously for 24 hours (mild injury), 48 hours (moderate injury), and 72 hours (severe injury). Lung specimens were taken for RNA, protein, and immunohistochemistry analysis (described below). All protocols were reviewed and approved by Yale University Institutional Animal Care and Use Committee.

For cell culture experiments, human type-II alveolar epithelial cells derived from induced pluripotent stem cells (AT2s^iPSC^) were used. AT2s^iPSC^ were generated using a “directed differentiation” approach, in which *in vivo* developmental milestones are recapitulated *in vitro* (46, 47). We used iPSCs containing a dual fluorochrome reporter for key genetic loci (*NKX2.1* identifies early lung progenitors; *SFTPC* is a marker for AT2s) (48). After iPSC differentiation through early differentiation stages to early lung progenitors, cells were seeded in extracellular culture matrix and allowed to differentiate into monolayered, epithelial spheres. AT2s^iPSC^ are primary cells viable in culture for several months (47). AT2s^iPSC^ exhibit characteristics of type-II distal respiratory epithelium. AT2s^iPSC^ were maintained in specific AT2 culture medium as described previously (49).

For *in vitro* hyperoxia exposure, cell cultures were exposed to hyperoxia (95% oxygen/5% carbon dioxide) in a tightly sealed modular exposure chamber (Billup-Rothberg, Del Mar, CA, USA; item #MIC-101) for pre-determined time points (39). For normoxia controls, cell cultures were maintained in an air-jacketed carbon dioxide (5%) incubator (VWR International, Radnor, PA, USA; model # 51014991). Cell supernatants were collected for lactate dehydrogenase (LDH) analysis. Cell lysates were collected for protein analysis.

### Cell Death Assays

For overall cell death, lactate dehydrogenase (LDH) was measured in cell culture supernatants using the Cytotoxicity Detection Kit (Roche Molecular Biochemicals, Indianapolis, IN, USA; item # 11644793001).

### Western Blotting

Specific proteins were evaluated and compared using Western blot analysis (50). For whole lung homogenates, lung tissue was homogenized in radioimmunoprecipitation assay (RIPA) buffer (Thermo Fisher Scientific, Waltham, MA, USA; catalog # 89900) and supplemented with protease inhibitor (Roche, Basel, Switzerland; reference # 11836170001) and phosphatase inhibitors sodium orthovanadate (New England Biolabs, Ipswich, MA, USA; item # P0758) and sodium fluoride (New England Biolabs; item # P0759). Western blotting was performed using a sodium dodecyl-sulfate polyacrylamide gel electrophoresis (SDS-PAGE) system (Bio-Rad, Hercules, CA, USA; item # 8658004). Samples were electrophoresed in a 4-20% polyacrylamide gel (Bio-Rad; item # 4568095) in tris-glycine-SDS buffer (Bio-Rad; catalog # 1610772) and transferred to a polyvinylidene fluoride (PVDF) membrane (Bio-Rad; item # 1704157). Samples were subsequently probed with rabbit anti-phospho-Beclin-1 (Ser93,-96) (1:1000 dilution; Thermo Fisher Scientific; catalog # PA5-67514), mouse anti-Beclin-1 (1:1000 dilution; Santa Cruz, Dallas, TX, USA; item # sc-48341), rabbit anti-LC3B (1:1000 dilution; Cell Signaling Technologies, Danvers, MA, USA; item # 2775), mouse anti-β-Actin (1:1000 dilution; Santa Cruz; item # sc-47778), and mouse anti-GAPDH (1:2500 dilution) primary antibodies overnight at 4°C. Samples probed with rabbit primary antibodies were subsequently probed with horseradish peroxidase (HRP)-linked anti-rabbit IgG (1:2500 dilution; Cell Signaling Technologies; item # 7074) secondary antibodies. Samples probed with mouse primary antibodies were subsequently probed with HRP-linked anti-mouse IgG (1:2500 dilution; Cell Signaling Technologies; item # 7076) secondary antibodies. Protein band densitometry analysis was completed using ImageJ software (version 1.53k; National Institutes of Health, USA). Full uncropped Western blots are provided as supplemental files (Supplemental File_Uncropped Western Blots).

### Reverse Transcription-Quantitative Polymerase Chain Reaction

Fold changes for specific mRNA were evaluated using reverse transcription-quantitative polymerase chain reaction (RT-qPCR) (51). Mouse lungs were collected and homogenized in TRIzol Reagent (Thermo Fisher Scientific, Carlsbad, CA, USA; item # 15596) and homogenized using the Precellys 24 bead mill homogenizer (Bertin Technologies, Montigny-le-Bretonneux, France). RNA was isolated and washed. RNase (Roche, Mannheim, Germany; item # 6011960) and DNAse (Roche; item # 04716728001) were added to the mixture. RNA integrity was confirmed, and RNA quantity was measured using a NanoDrop 2000 microvolume spectrophotometer (Thermo Fisher Scientific, Wilmington, DE, USA; item # ND-2000). 1 µg RNA was synthesized to cDNA by reverse-transcription using the iScript cDNA Synthesis Kit (Bio-Rad, Hercules, CA, USA; item # 1708891). Real-time PCR was performed in 10 µl reactions containing 5 µl SsoFast EvaGreen Supermix with Low ROX (Bio-Rad; item # 1725211) diluted 1:2, 2 µl cDNA template (diluted 1:10), 0.5 µl forward primer, 0.5 µl reverse primer, and 2 µl deionized water. Quantitative PCR was performed with the ViiA 7 Real-Time PCR System (Applied Biosystems, Waltham, MA). Primers for real-time PCR were manufactured at the Keck Biotechnology Research Laboratory at Yale University (New Haven, CT, USA). The following primers were used: BCN1, 5′-CTGAAACTGGACACGAGCTTCAAG-3′ and 5′-CCAGAACAGTATAACGGCAACTCC-3′ (52); BCN1, ′-TTT TCT GGA CTG TGT GCA GC-3′ and 5′-GCT TTT GTC CAC TGC TCC TC-3′ (53); Atg5, 5′-GACAAAGATGTGCTTCGAGATGTG-3′ and 5′-GTAGCTCAGATGCTCGCTCAG-3′ (52); ATG7, 5′-ATGCCAGGACACCCTGTGAACTTC-3′ and 5′-ACATCATTGCAGAAGTAGCAGCCA-3′ (54); LC3a,5′-AGCTTCGCCGACCGCTGTAAG-3′ and 5′-CTTCTCCTGTTCATAGATGTCAGC-3′ (52); and LC3b, 5′-CGGAGCTTTGAACAAAGAGTG-3′ and 5′-TCTCTCACTCTCGTACACTTC-3′ (52). The levels of mRNA were normalized to glyceraldehyde-3-phosphate dehydrogenase (GAPDH). Fold changes were calculated using the 2^-ΔΔCT^ method.

### Immunohistochemistry

Mice were anesthetized and euthanized, the pulmonary intravascular space was washed until clear, and whole lungs were excised from mice and immediately placed in 4°C buffered formalin overnight for paraffin embedding (44). Immunohistochemistry was performed on 5-µm sections by the Yale Histopathology Core. Antigen retrieval was completed in antigen retrieval buffer. Lung sections were labelled with rabbit anti-Beclin-1 (Abcam; catalog # ab62557) antibodies, 1:400 overnight at 4°C. Subsequently, sections were incubated in anti-rabbit secondary antibodies conjugated to horseradish peroxidase. Slides were analyzed with an Eclipse Ni upright microscope (Nikon, Melville, NY, USA). Images were captured using a DS Ri2 microscope camera (Nikon) with NIS Elements software version 4.30 (Nikon). Cell-specific localization of BCN1-positive cells was determined by two independent observers (Drs. Harris and Zhang). Observers were blinded to each sample’s allocation of experimental group (i.e., normoxia vs. hyperoxia). H-scores were calculated for BCN1 by assigning a staining intensity ordinal value for each cell as follows: 0 for no signal, 1 for mild signal, 2 for moderate signal, and 3 for strong signal (55). The percentage of cells for each staining intensity ordinal value was multiplied by the respective staining intensity ordinal value, and the H-score was calculated by adding the four products together, such that H-score = 0(% cells counted as none) + 1(% cells counted as mild) + 2(% cells counted as moderate) + 3 (% cells counted as strong). An H-score ranging from 0-300 was generated. 100 cells were counted for each sample.

### Statistical Analysis

Molecular data were examined by analyses of variance to estimate means and standard errors. Statistical significance was assessed by two-tailed *t*-tests for parametric data and Mann Whitney U tests for non-parametric data. Survival data were examined by constructing Kaplan-Meier plots and statistical significance assessed by log-rank chi square tests. Graphs and statistical analyses were completed using GraphPad Prism version 10.1.1 (GraphPad Software version 10.1.1, La Jolla, CA, USA). Standard deviations are reported in all graphs to show variability of the data. In addition, precise *P* values are reported in all graphs.

## RESULTS

### BCN1 is increased in the lung *in vivo* in HALI

The effects of HALI on BCN1 were examined in mouse lungs. HALI was induced in wildtype (WT) mice by exposure to 100% oxygen for 24, 48, and 72 hours, and lungs were analyzed for BCN1. Whole-lung BCN1 [p-and total-fraction] was evaluated via Western blot (Figure 1A-C). Phosphorylation of BCN1 was evaluated for the serine residue 91 and 94 (Ser-91,-94) for mouse or Ser-93 and-96 for human, which leads to activation of the BCN1-VPS34 complex and subsequent initiation of autophagy (56). In HALI, the ratio of p-/total BCN1 increased at 48 hours compared with 24 hours, however no differences were observed between normoxia and 48 hours (Figure 1A-B). The p-BCN1/β-Actin ratio increased at 48 hours compared with normoxia and 24 hours (Figure 1A, 1C). The total BCN1/β-Actin ratio increased at 48 hours compared with normoxia and 24 hours (Figure 1A, 1D). In addition, whole-lung expression of BCN1 was evaluated via reverse transcription-quantitative polymerase chain reaction (RT-qPCR; Figure 1E). In HALI, BCN1 expression increased at 48 and 72 hours compared with normoxia (Figure 1E). To confirm these findings, we used an additional BCN1 primer for RT-qPCR, and the results were similar that BCN1 expression increased in HALI (Supplemental Figure S1A). In addition, BCN1 in the lung in HALI was also evaluated via immunohistochemistry (Figure 1F). H-scores, an immunohistochemical grading system used to quantify the intensity of immunohistochemical staining (55), were calculated for each sample. In HALI, BCN1 increased in alveolar epithelial cells at 48 hours compared with normoxia (Figure 1F). Together, these data indicate that, in lungs of WT mice in HALI, both p-BCN1 and total BCN1 increase at 48 hours compared with normoxia. Although the BCN1/β-Actin increases at 48 hours compared with normoxia, the p-/total BCN1 ratio does not change. These data also indicate that BCN1 increases significantly in the alveolar epithelium at 48 hours in HALI.

**Figure 1.**
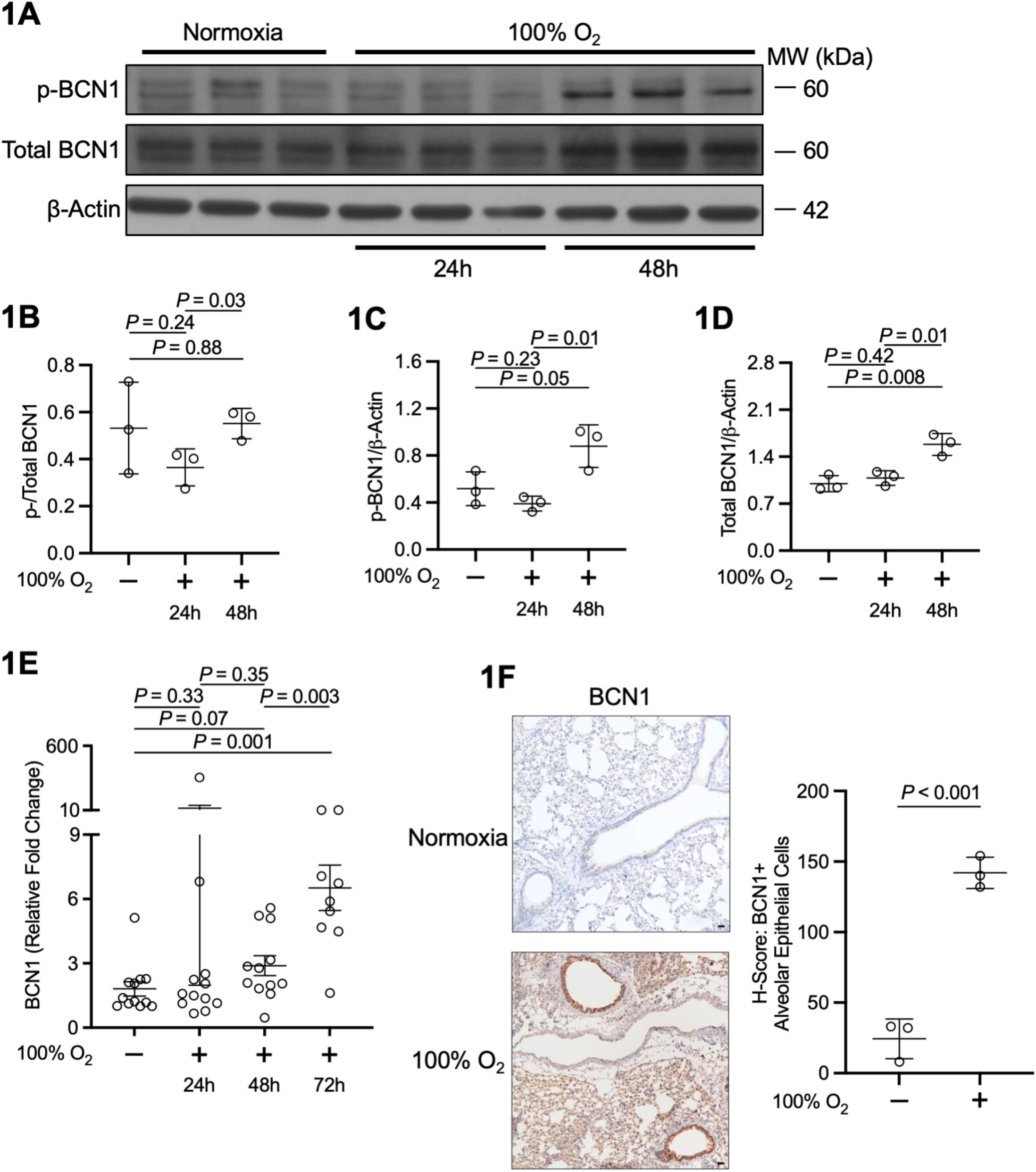
Pulmonary BCN1 is increased in HALI. (A-E) Effects of HALI (100% oxygen) on BCN1 in the lungs of WT mice at 24, 48, and 72 hours compared with normoxia (*N* = 4-6 mice/group, repeated once). (A) Representative Western blot on whole lungs for BCN1 (p-and total) at 24 and 48 hours compared with normoxia. (B-C) Quantitative densitometry for (B) p-/total BCN1, (C) p-BCN1/β-Actin, and (D) total BCN1/β-Actin is shown. Unpaired t test. (E) RT-qPCR on whole lungs for BCN1 at 24, 48, and 72 hours. Unpaired t test. (F) Immunohistochemistry on lungs for BCN1 at 48 hours compared with normoxia. Quantification of H-score for alveolar epithelial cells is shown. Unpaired t test. BCN1, Beclin-1.

### HALI leads to alterations in autophagy markers in the lung *in vivo*

Because BCN1 modulates autophagy, the effects of HALI on autophagic markers were examined in mouse lungs. WT mice were exposed to 100% oxygen for 24, 48, and 72 hours to induce HALI, and lungs were analyzed for markers of autophagy. Whole-lung LC3B was evaluated via Western blot (Figure 2A-C). Endogenous LC3B-I is converted into LC3B-II [the phosphatidylethanolamine (PE)-conjugated form], and LC3B-II associates with autophagosome membranes (57). The LC3B-II/β-Actin ratio was measured, as LC3B-II levels correlate well with the number of autophagosomes (57). Whole-lung LC3B-II levels did not change at 24, 48, and 72 hours in HALI compared with normoxia (Figure 2A-B). Whole-lung LC3B-II levels were increased significantly at 48 hours compared with 24 hours (Figure 2A-B), suggesting increased autophagosomes. Next, whole-lung LC3B-II/-I ratios were assessed via Western blot (Figure 2A, 2C). LC3B-II/-I ratios were decreased at 24 and 72 hours in HALI compared with normoxia (Figure 2A, 2C), suggesting decreased autophagic flux (i.e., decreased level of cellular autophagic activity) (57) in the lung at those time points. No differences were observed in LC3B-II/-I ratios at 48 hours compared with normoxia (Figure 2A, 2C). In addition, whole-lung expression of the autophagy genes autophagy related 5 (ATG5), ATG7, LC3A, and LC3B were examined in mouse lungs via RT-qPCR (Figure 2D-E, Supplemental Figure S2A). ATG7 expression decreased at 72 hours in HALI compared with normoxia (Figure 2D). There was also decreased ATG7 at 72 hours compared with 48 hours (Figure 2D). In addition, LC3A expression decreased at 48 and 72 hours in HALI compared with normoxia (Figure 2E). However, ATG5 expression increased at 24 and 48 hours in HALI compared with normoxia (Supplemental Figure 2A). In addition, there was increased LC3B expression at 48 and 72 hours compared with normoxia (Supplemental Figure 2B). Together, these data suggest that, in lungs of WT mice in HALI, there are significant alterations in autophagic markers and autophagic flux decreases at 24 and 72 hours. Together with the analyses of BCN1, these data indicate that, in the lung of WT mice in HALI, BCN1 increases while autophagic flux is reduced.

**Figure 2.**
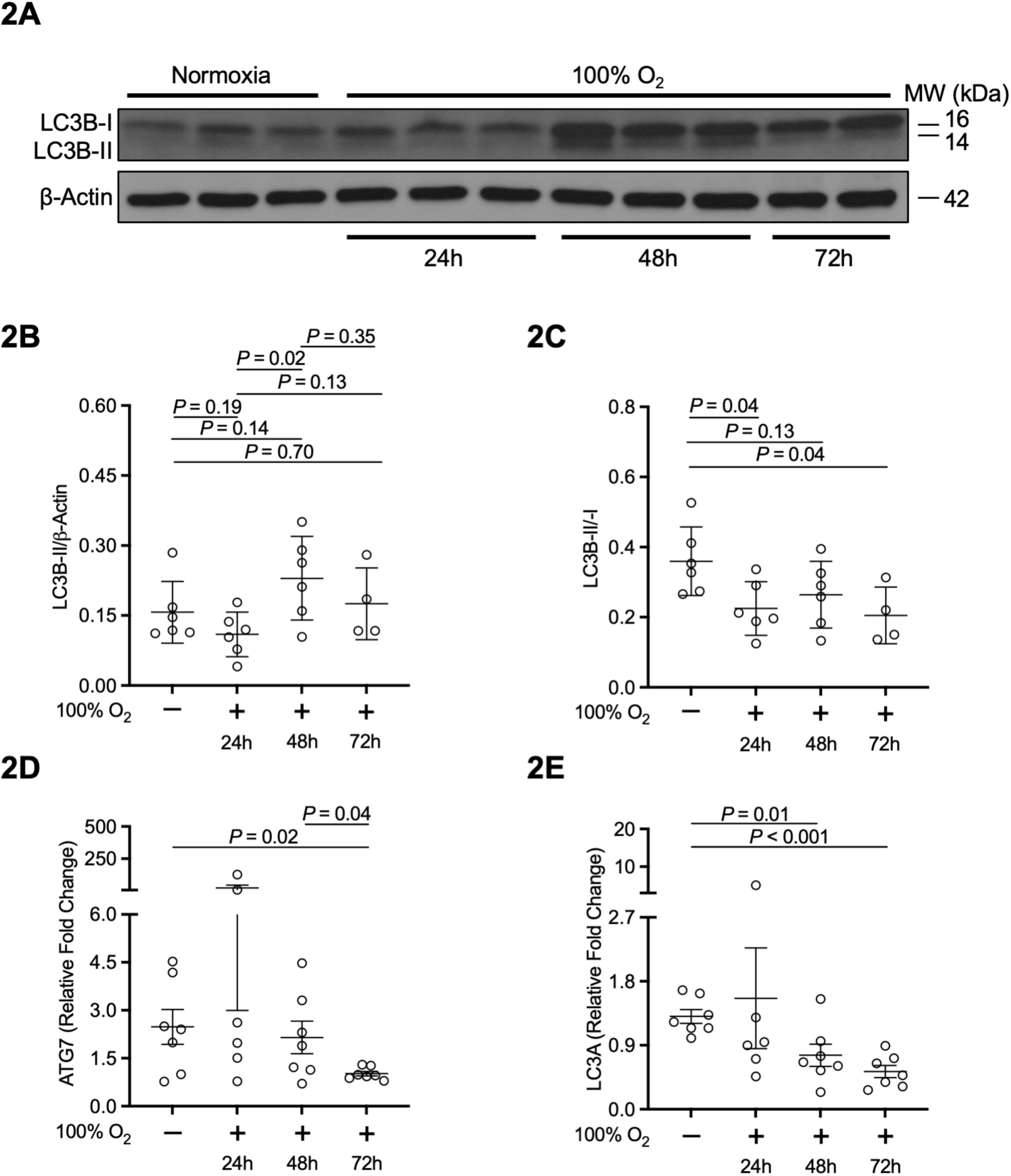
HALI exerts significant alterations on autophagy markers in the lung *in vivo*. (A-E) Effects of HALI (100% oxygen) on autophagic markers in the lungs of WT mice at 24, 48, and 72 hours compared with normoxia (*N* = 4-6 mice/group, repeated once). (A) Representative Western blot on whole lungs for LC3B. (B-C) Quantitative densitometry for (B) LC3B-II/β-Actin and (C) LC3B-II/-I is shown. Unpaired t test. (D-E) RT-qPCR on whole lungs for (D) ATG7 and (E) LC3A. Unpaired t test. ATG7, autophagy related 7. LC3A, microtubule-associated protein 1A-light chain.

### HALI leads to increased LDH release and decreased LC3B-II/-I ratios in human AT2s^iPSC^

Based on our results of BCN1 modulation in lung epithelium (Figure 1F) and the well-established effects of HALI on the alveolar epithelium, effects of HALI on LC3B were examined in human AT2s^iPSC^. AT2s^iPSC^ in culture formed organoids (alveolospheres) and expressed the markers NKX2.1 and SPC that are associated with type-II alveolar epithelial cells (Figure 3A). AT2s^iPSC^ were exposed to 95% oxygen to induce HALI and compared to normoxia for 48 and 72 hours in the presence of DMSO (10 µM). Cell supernatants were analyzed for lactate dehydrogenase (LDH), a marker of cell death. Cell lysates were analyzed for LC3B-II/-I ratios via Western blot to determine autophagic flux (57). HALI led to increased LDH release in AT2s^iPSC^ compared with normoxia at 72 hours (Figure 3B), indicative of increased cell death. LC3B-II/-I ratios via Western blot were decreased at 48 hours in HALI compared with normoxia (Figure 3C), suggesting decreased autophagic flux. These results were consistent with our *in vivo* observations that HALI leads to reduced LC3B-II/-I ratios.

**Figure 3.**
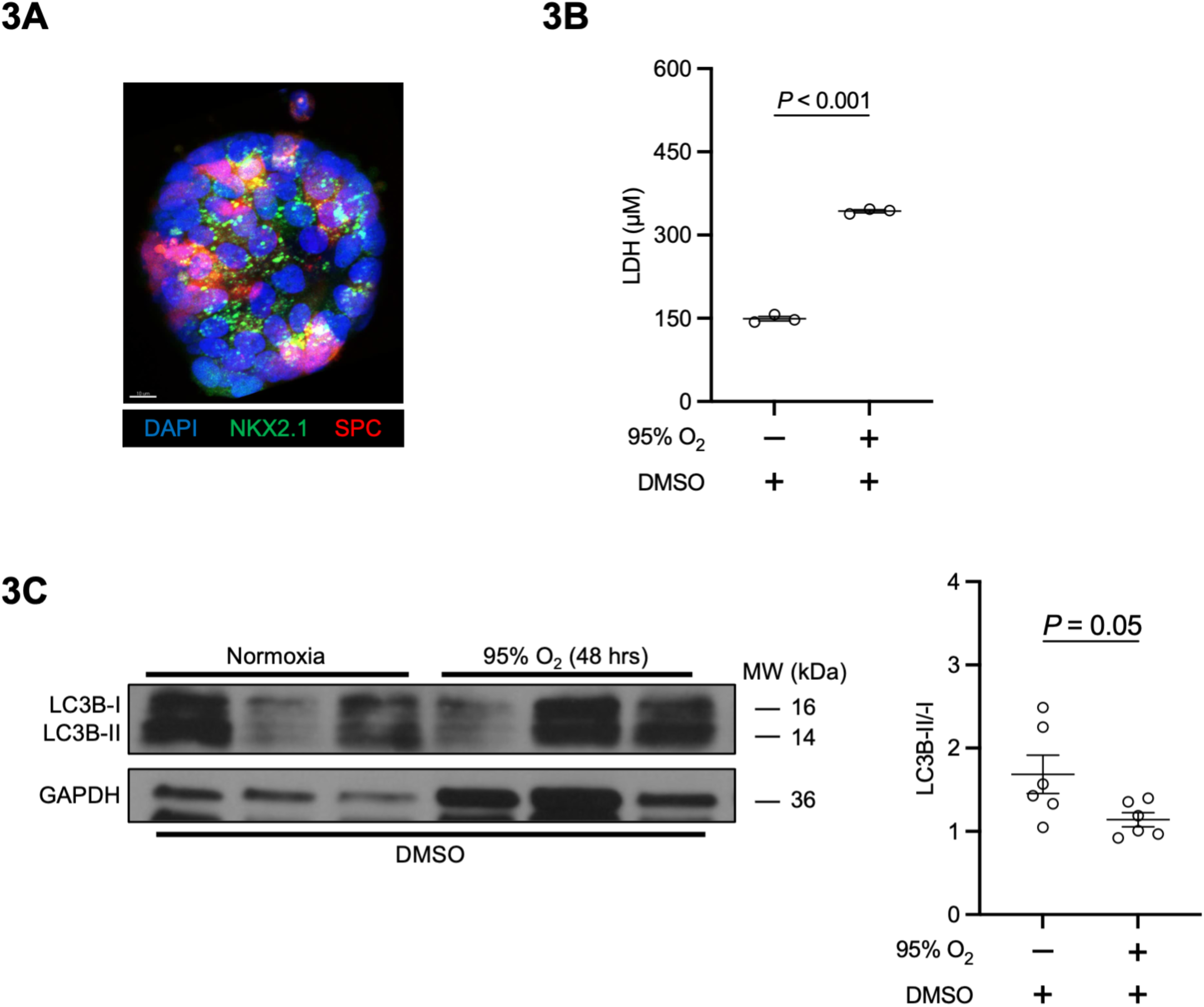
HALI induces increased LDH release and reduced LC3-II/-I ratios in AT2s^iPSC^. (A-C) Effects of HALI (95% oxygen) on LDH, a marker of cell death, and LC3B, a marker for autophagy, were examined in AT2s^iPSC^ in the presence of DMSO (10 µM) and compared with normoxia. (A) SPC+/NKX2.1+ alveolospheres. Images taken on a Zeiss LSM880 Airyscanner confocal microscope. Scale bar = 10 µM. SPC-producing cells are identified with orange arrows. (B) Supernatant LDH, 72 hours. Unpaired t test. (C) Representative Western blot on cell lysates for LC3B with quantitative densitometry for LC3B-II/-I. Unpaired t test. DAPI, 4’,6-diamidino-2phenylindole. DMSO, dimethylsulfoxide. GAPDH, glyceraldehyde-3-phosphate dehydrogenase. LDH, lactate dehydrogenase. LC3B, microtubule-associated protein 1B-light chain. SPC, surfactant protein C.

### Genetic EGFR inhibition regulates BCN1 in the lung *in vivo* in HALI

We previously showed that EGFR^Wa5/+^ mice, which contain a point mutation that decreases EGFR activity significantly (43), have improved survival, decreased ALI, and decreased alveolar epithelial apoptotic cell death in HALI compared with WT (39). Based on studies showing regulation of BCN1 by EGFR (42), we hypothesized that BCN1 was altered in EGFR^Wa5/+^ mice in HALI compared with WT and that BCN1 regulation might provide a mechanism that provides protection in EGFR^Wa5/+^ mice in HALI. Effects of HALI (100% oxygen) on BCN1 in the lung were examined in EGFR^Wa5/+^ and WT mice compared with normoxia at 24, 48, and 72 hours. BCN1 [p-(Ser-93,-96) and total-fraction] was evaluated via Western blot (Figure 4A-B). In normoxia, there were no differences in p-/total BCN1 ratios and total BCN1 in EGFR^Wa5/+^ mice compared with WT (Figure 4A). In HALI at 48 hours, EGFR^Wa5/+^ mice showed decreased p-/total BCN1 ratios and increased total BCN1 in the lung compared with WT (Figure 4B). Together, these data indicate that, in HALI in the lung, EGFR^Wa5/+^ mice have decreased whole-lung p-/total BCN1 ratios and increased total BCN1 compared with WT.

**Figure 4.**
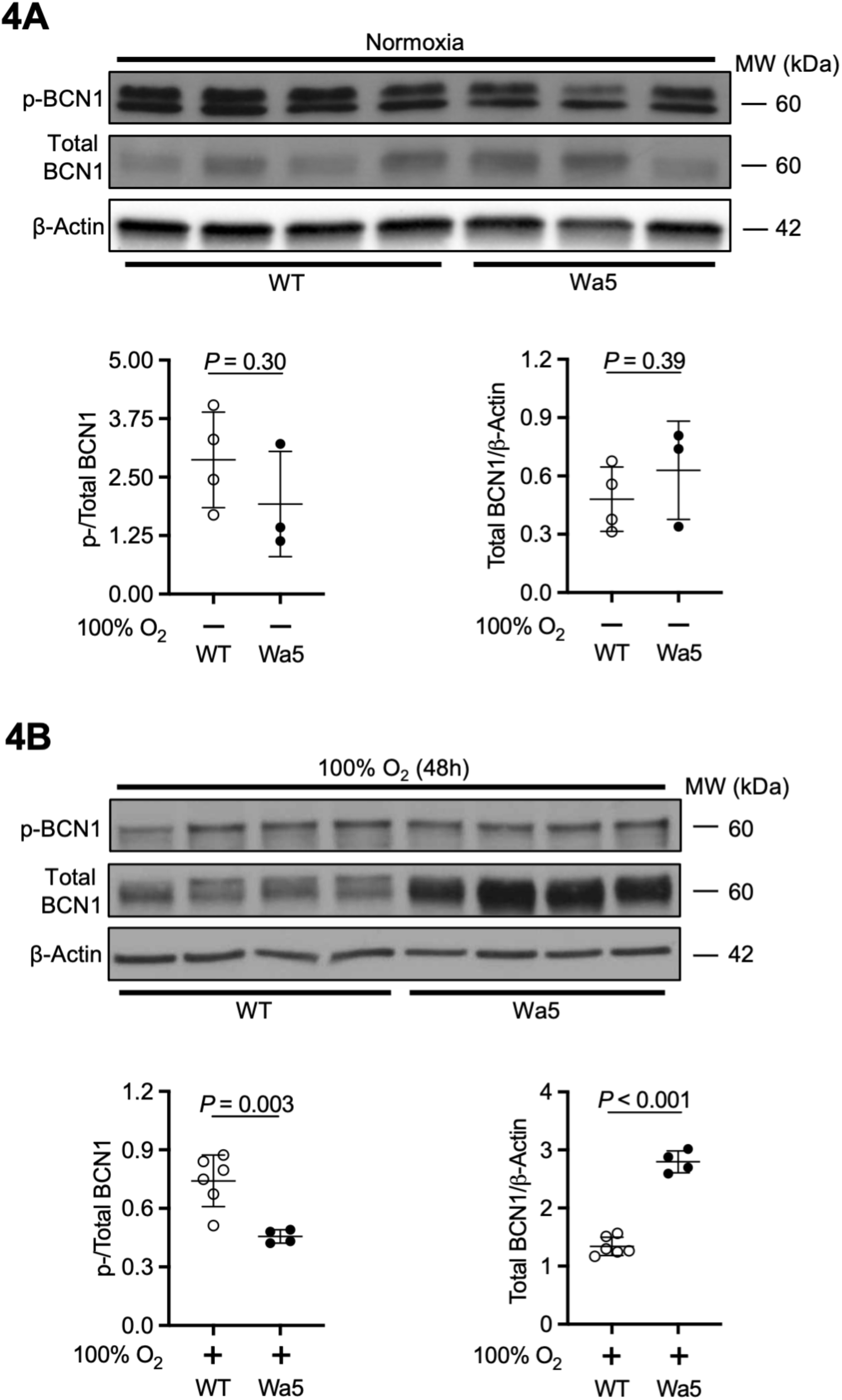
Genetic EGFR inhibition regulates BCN1 in the lung *in vivo* in HALI. (A-D) Effects of HALI (100%) oxygen on BCN1 in the lungs of EGFR^Wa5/+^ and WT mice compared with normoxia (*N* = 4-6 mice/group, repeated twice). (A-B) Representative Western blot for BCN1 (p-and total) on whole lungs with quantitative densitometry for p-/total BCN1 and total BCN1/β-Actin. (A) Normoxia. (B) 100% oxygen, 48 hours. Unpaired t test. BCN1, Beclin-1. Wa5, EGFR^Wa5/+^.

### Genetic EGFR inhibition results in decreased LC3B-II/I ratios in the lung *in vivo* in HALI

Because BCN1 phosphorylation at Ser93,-96 leads to autophagosome biogenesis (56), we hypothesized that autophagy would be affected in the lungs of EGFR^Wa5/+^ mice compared with WT in HALI. Effects of HALI (100% oxygen) on whole-lung autophagic markers were examined in EGFR^Wa5/+^ and WT mice and compared with normoxia. LC3B was assessed via Western blot. LC3B-II/β-Actin ratios, which correlate with autophagosome number, were evaluated. In addition, LC3B-II/I ratios, which are suggestive of autophagic flux activity, were evaluated. No differences were observed in LC3B-II/β-Actin and LC3B-II/-I ratios in the lung in EGFR^Wa5/+^ and WT mice in normoxia (Figure 5A). In HALI at 48 hours, LC3B-II/β-Actin and LC3B-II/-I ratios were both decreased in the lung in EGFR^Wa5/+^ mice compared with WT (Figure 5B). These data suggest that there is decreased autophagosome number and decreased autophagic flux in the lungs of EGFR^Wa5/+^ mice compared with WT in HALI.

**Figure 5.**
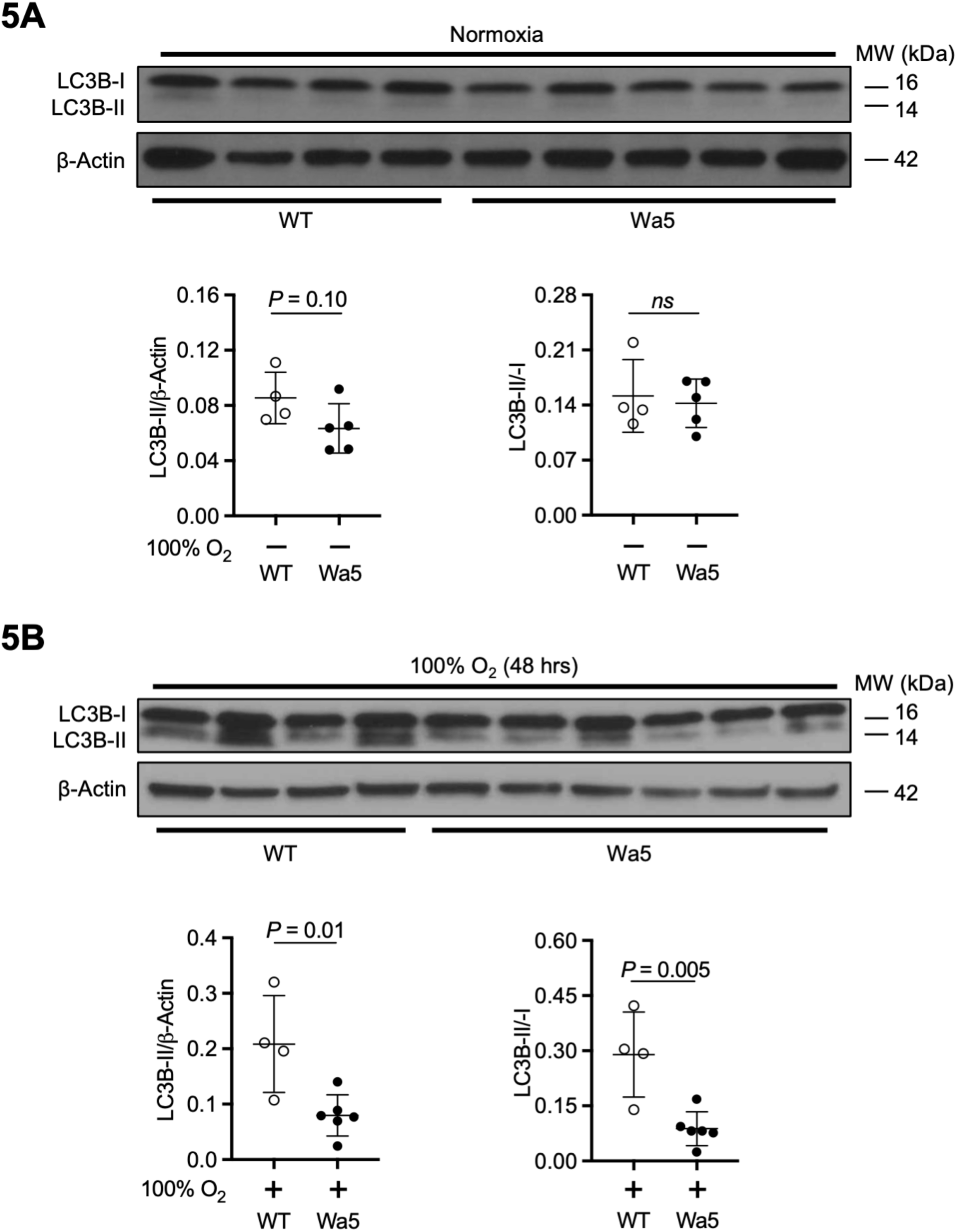
Genetic EGFR inhibition leads to reduced LC3B-II/-I ratios in the lung *in vivo* in HALI. (A-C) Effects of HALI (100% oxygen) on autophagy in the lungs of EGFR^Wa5/+^ and WT mice compared with normoxia (*N* = 4-6 mice/group, repeated twice). (A-B) Representative Western blot on whole lungs for LC3B with quantitative densitometry for LC3B-II/β-Actin and LC3B-II/-I. (A) Normoxia. (B) 100% oxygen, 48 hours. Unpaired t test. LC3B, microtubule-associated protein 1B-light chain. Wa5, EGFR^Wa5/+^.

### Administration of wortmannin worsens survival in HALI *in vivo*

Despite studies showing a beneficial role of autophagy in HALI (18), the role of BCN1-mediated autophagy in HALI is not well defined. To investigate the role of BCN1-mediated autophagy in HALI, we used the pharmacologic agent wortmannin, a mold metabolite that inhibits PI3K and exerts persistent inhibitory effects on the VPS34 complex (45). Effects of wortmannin (1 mg/kg administered intraperitoneally) on survival in HALI (100% oxygen) were examined in WT mice and compared with mice treated with vehicle control. Wortmannin significantly worsened survival in HALI (Figure 6A), suggesting that inhibition of BCN1-mediated autophagy is deleterious in HALI.

**Figure 6.**
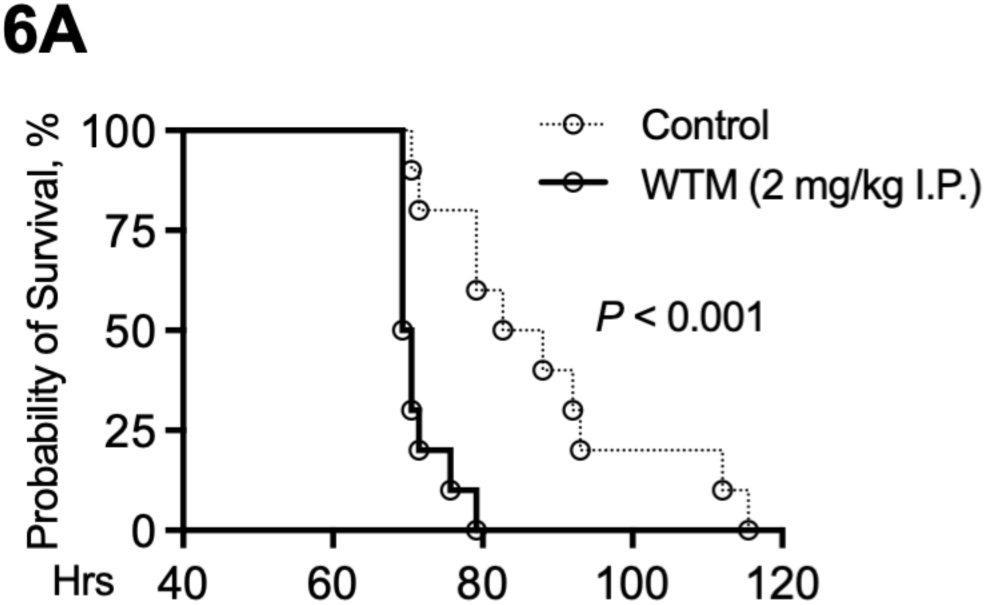
Wortmannin administration worsens survival in HALI. (A) Effects of wortmannin (2 mg/kg intraperitoneal) on survival in WT mice in HALI (100% oxygen) compared with vehicle control. Log-rank test. WTM, wortmannin.

## DISCUSSION

Approximately 180,000 patients are affected with ARDS each year in the United States, and mortality for the condition is as high as 40% (22). There are no medical therapies for ARDS. In addition, ARDS was the leading cause of ICU admission during the COVID pandemic (58, 59). Thus, it is critical to develop therapeutics for ARDS. Because HALI is a form of ARDS induced purely by oxidative damage (11) and, in animals, is a model for “sterile” (non-infectious) ARDS (60), elucidation of molecular pathways in HALI enables identification of cell death pathways that are active in ARDS mediated by oxidative damage. Thus, therapeutic targets identified in HALI have the potential to be candidate therapeutic targets for ARDS, especially as our ability to identify patient-specific phenotypes in ARDS improves.

This study shows that pulmonary BCN1 is increased in HALI. In addition, HALI led to significant alterations in autophagy markers in the lung *in vivo*, suggesting decreased autophagic flux. In addition, HALI led to increased LDH release and reduced LC3B-II/-I ratios in human AT2s^iPSC^ *in vitro*. We previously showed EGFR^Wa5^ mice with genetically reduced EGFR activity have improved survival, reduced ALI, and decreased alveolar epithelial apoptotic cell death in HALI (39). In the current study, these mice contained decreased p-/total BCN1 ratios, increased total BCN1, and reduced LC3-II/-I ratios in the lung in HALI. *In vivo*, administration of the PI3K inhibitor wortmannin, which inhibits BCN1-mediated autophagy, increased mortality in HALI. These findings suggest a key role of BCN1-mediated autophagy in HALI that is regulated by EGFR. Regulation of BCN1 and autophagy by EGFR and its role in HALI warrants further investigation as a mechanism that could be therapeutically targeted in ALI/ARDS.

Our findings are consistent with previous studies showing hyperoxia-mediated effects on autophagy (18). In addition, our findings compliment previous studies which link EGFR activation and autophagy. EGFR activation is known to regulate the mammalian target of rapamycin (mTOR) pathway via PI3K/v-akt murine thymoma viral oncogene homolog 1 (AKT), and this pathway is known to regulate autophagy (61). In addition, EGFR is known to regulate BCN1 through multiple mechanisms in other cellular conditions. EGFR inhibits autophagy in cancer cells by binding to BCN1, leading to multisite tyrosine phosphorylation, enhanced binding of the BCN1 complex to inhibitors, and subsequent reduced class III PI3K (i.e., VPS34 complex) activity (42). However, serum starvation induces arrest of inactive EGFR at the endosome, which then binds to Run domain BCN1 interacting and cysteine-rich containing protein (Rubicon), releasing the BCN1 complex for autophagy initiation (41, 62). While our studies in HALI implicate EGFR regulation of a specific serine phosphorylation site of BCN1 associated with autophagy initiation, serine residue 91 and 94 (Ser-91,-94) for mouse or Ser-93,-96 for human, the molecular machinery involved in EGFR regulation of BCN1 in HALI remains to be elucidated.

Our results suggest that BCN1-mediated autophagy is a key molecular pathway in HALI, however critical questions remain. The role of BCN1 and autophagy in HALI is not well defined (63, 64), and the role of BCN1 and autophagy in ALI is an area of active investigation (65–68). The results of the current study allow us to speculate that BCN1 and autophagy, which are regulated by EGFR, modulate cell death in HALI. Whether regulated cell death (RCD) induced by autophagy, called autophagy-dependent cell death (ADCD) (69, 70), plays a significant role in HALI is not clear and warrants further study. In addition, increased BCN1 levels can bind with and sequester proteins in the BCL2 family and consequently upregulate apoptotic cell death while simultaneously downregulating autophagy, a mechanism which has been shown to be clinically relevant in cardiac disease (71). How EGFR regulates the interaction of BCN1 with BCL2 family proteins in HALI is unclear and warrants further study.

## LIMITATIONS

This study has some limitations. First, the data confirming that EGFR^Wa5/+^ mice are protected in HALI and have reduced alveolar epithelial cell death that we previously showed (39) are not included in this manuscript. This is relevant because the current study builds on these data and provides a potential mechanism which might relate to increased survival and reduced apoptotic cell death in EGFR^Wa5/+^ mice in HALI. Whether alterations in BCN1 and autophagy contribute to decreased apoptotic cell death in in EGFR^Wa5/+^ mice in HALI warrants further study but is outside the scope of this manuscript.

Second, administration of wortmannin, a PI3K inhibitor which blocks BCN1-mediated autophagy, led to worsened survival in HALI. However, wortmannin also transiently inhibits class I PI3K and exerts effects on phylogenetically related kinases such as mTOR, DNA-dependent protein kinase (DNA-PK), and ataxia telangiectasia mutated (ATM) protein kinase (45). Our results show that wortmannin administration is deleterious in HALI, however whether wortmannin’s deleterious effects in HALI are from BCN1-mediated autophagy inhibition in the lung or from a separate mechanism remains to be elucidated. Genetic techniques targeting BCN1 (72) and pharmacologic agents that target specific PI3K isoforms (45) can help further elucidate the role of BCN1-mediated autophagy in HALI, but these studies are outside the scope of this manuscript.

Finally, our data demonstrate an increase in alveolar epithelial BCN1 in HALI. However, EGFR regulation of BCN1 activation specific to lung epithelium in HALI warrants further investigation, as additional cell types may be involved. BCN1 has been shown to be key to regulation of inflammation in myeloid cells (73). BCN1 has also been shown to play an important role in endothelial cell inflammation and barrier disruption (74). Cell-specific conditional knockout approaches could help further elucidate the mechanisms of EGFR regulation of BCN1 specific to the lung epithelium in HALI, but these studies are outside the scope of this manuscript.

## CONCLUSIONS

HALI resulted in increased BCN1 in mouse lung alveolar epithelium *in vivo*. HALI led to significant alterations in pulmonary autophagy markers, including reduced LC3B-II/-I ratios, suggesting decreased autophagic flux. *In vitro* in human AT2s^iPSC^, HALI led to increased LDH release and reduced LC3B-II/-I ratios, suggesting decreased autophagic flux. *In vivo* in HALI, EGFR^Wa5/+^ mice, which are protected in HALI and contain decreased alveolar epithelial apoptosis, had decreased p-/total BCN1 ratios, increased total BCN1, and reduced LC3B-II/-I ratios in the lung. Administration of wortmannin, a PI3K inhibitor which blocks BCN1-mediated autophagy, worsened survival in HALI. Regulation of BCN1 by EGFR is a novel mechanism in HALI with therapeutic potential that warrants further study.

## ETHICS, CONSENT TO PARTICIPATE, AND CONSENT TO PUBLISH DECLARATIONS

Not applicable.

## DATA AVAILABILITY

The data that supports the findings of this study are available in the supplementary material of this article. The data that support the findings of this study are available on request from the corresponding author.

## PREPRINT

A preprint has previously been published Harris *et al.*, 2024 (75).

## ACKNOWLEDGMENTS

The authors would like to thank Rob Homer for his time, generosity and contributions to this manuscript. The preprint for this manuscript is available at https://www.biorxiv.org/content/10.1101/2024.05.24.595621v2.

## AUTHOR CONTRIBUTIONS

ZMH, AK, LS, MS, GR, HW, MG, MJK, and JLK conceived and designed research. ZMH, AK, JK, EPM, GS, YS, BH, HJS, JJ, BC, LP, DU, AM, and LS performed experiments. ZMH, AK, KM, BH, HJS, LS, and XZ analyzed data. ZMH, AK, LS, MS, GR, HW, MJK, and JLK interpreted results of experiments. ZMH prepared figures. ZMH drafted manuscript. ZMH, GR, HW, MJK, and JLK edited and revised manuscript. All authors approved final version of manuscript.

## PATIENT AND PUBLIC INVOLVEMENT STATEMENT

Patients/public were not involved in the research.

**Supplemental Figure S1. HALI leads to increased BCN1 in the lung *in vivo*.** (A) Effects of HALI (100% oxygen) on BCN1 in the lungs of WT mice at 24, 48, and 72 hours compared with normoxia (*N* = 4-6 mice/group, repeated once). RT-qPCR on whole lungs for BCN1. Unpaired t test. BCN1, Beclin-1.

**Supplemental Figure S2. HALI regulates autophagy markers in the lung *in vivo*.** (A-B) Effects of HALI (100% oxygen) on autophagy markers in the lungs of WT mice at 24, 48, and 72 hours compared with normoxia (*N* = 4-6 mice/group, repeated once). RT-qPCR on whole lungs for (A) ATG5 and (B) LC3B. Unpaired t test. ATG5, autophagy related 5. LC3B, microtubule-associated protein 1B-light chain.

Full uncropped Western blots are provided as supplemental files (Supplemental File_Uncropped Western Blots).

