## Supplementary figures and images for "Epidermal Growth Factor Receptor Regulates Beclin-1 in Hyperoxic Acute Lung Injury"

### Supplementary Figure 1

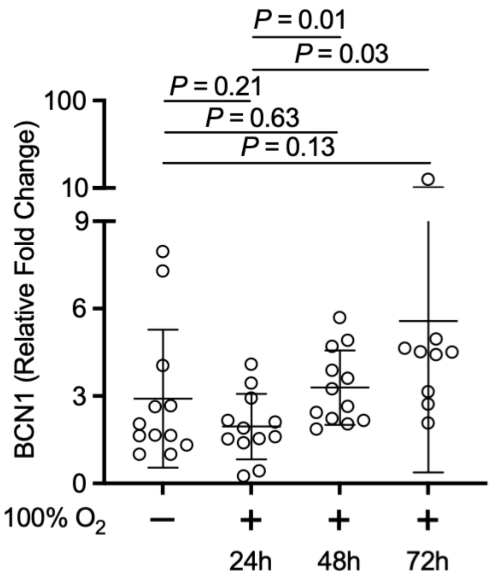

### Supplementary Figure 2

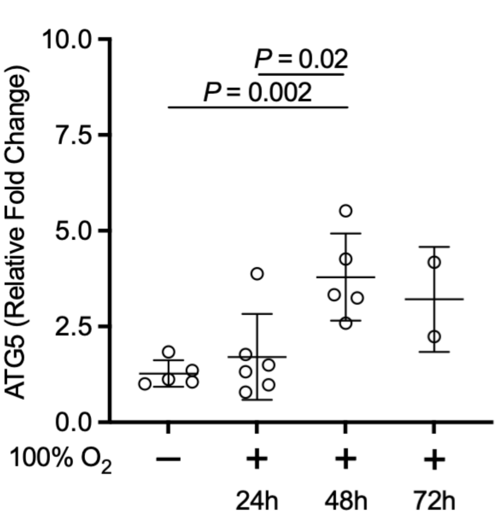

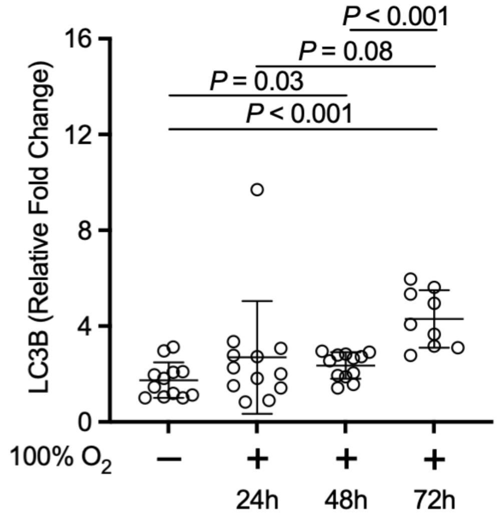
